# Revealing the evolutionary history of a reticulate polyploid complex in the genus *Isoëtes*

**DOI:** 10.1101/2020.11.04.363374

**Authors:** Jacob S. Suissa, Sylvia P. Kinosian, Peter W. Schafran, Jay F. Bolin, W. Carl Taylor, Elizabeth A. Zimmer

## Abstract

- Polyploidy and hybridization are important processes in the evolution of spore-dispersed plants. Few studies, however, focus these dynamics in heterosporous lycophytes, such as *Isoëtes*, where polyploid hybrids are common and thought to be important in the generation of their extant diversity. We investigate reticulate evolution in a complex of western North American quillworts (*Isoëtes*) and provide insights into the evolutionary history of hybrids, and the role of polyploidy in maintaining novel diversity.
- We utilize low copy nuclear markers, whole plastomes, restriction site-associated DNA sequencing, cytology, and reproductive status (fertile or sterile) to investigate the reticulate evolutionary history of western North American *Isoëtes*.
- We reconstruct the reticulate evolutionary history and directionality of hybridization events in this complex. The presence of high level polyploids, plus frequent homoploid and interploid hybridization suggests that there are low prezygotic reproductive barriers in this complex, hybridization is common and bidirectional between similar—but not divergent—cytotypes, and that allopolyploidization is important to restore fertility in some hybrid taxa.
- Our data provide five lines of evidence suggesting that hybridization and polyploidy can occur with frequency in the genus, and these evolutionary processes may be important in shaping extant *Isoëtes* diversity.

## Introduction

Processes such as hybridization and genome duplication (polyploidization) can lead to evolutionary histories that are better explained through reticulating, rather than bifurcating, branches on a phylogenetic tree (Soltis & Soltis, 2000; Linder & Rieseberg, 2004; Nakhleh *et al.*, 2005; Degnan & Rosenberg, 2009; Edelman *et al.*, 2019). These evolutionary processes are important because they lead to the generation and maintenance of novel lineages. These lineages form the basis for reticulate species complexes, which are webs of related taxa involving multiple parental species, sterile (or partially sterile) interspecific hybrids (Dobzhansky, 1982), and fertile auto- or allopolyploids (i.e., products of genome duplication with one or more prenatal genomes, respectively; Stebbins, 1969; Grant, 1971; Rieseberg, 1997). Sterile hybrids within these complexes are sometimes considered evolutionary dead ends, but through allopolyploidization, fertility can be restored because each chromosome has a match in its newly duplicated genome (Stebbins, 1947; Wagner & Wagner, 1980; Soltis & Soltis, 2000). In addition, allopolyploidy can lead to fixed genomic variation (heterozygosity) allowing for lower inbreeding depression following self-fertilization (Soltis & Soltis, 2000; Husband *et al.*, 2008; Soltis *et al.*, 2014; Haufler *et al.*, 2016). For these reasons, allopolyploidy is suggested to be evolutionarily advantageous within some lineages—however the importance of this process as a broader driver of diversification is debated (Mayrose *et al.*, 2011, 2015; Soltis *et al.*, 2014; Landis *et al.*, 2018).

Reticulate species complexes are common in plants (Burnier *et al.*, 2009; Nauheimer *et al.*, 2019; Sandstedt *et al.*, 2020), especially in spore-dispersed vascular plants (ferns and lycophytes) (Barrington *et al.*, 1989; Wood *et al.*, 2009; Sigel, 2016). Among spore dispersed plants, a majority of this research focuses on homosporous ferns, with less work on the lycophytes, and especially the genus *Isoëtes* L (Isoetaceae) where these processes are thought to be rampant (Taylor & Luebke, 1988; Hickey *et al.*, 1989; Britton & Brunton, 1989; Taylor & Hickey, 1992; Brunton & Britton, 2000; Small & Hickey, 2001; Pereira *et al.*, 2018). *Isoëtes* is a cosmopolitan genus comprising 250-350 species, ~60% of which are polyploid (Troia *et al.*, 2016). This genus has been described as a “living fossil” as it is the last remaining lineage of the ancient Isoetalean lycopsids—a clade that dates back to the middle Devonian (~400mya; Pigg, 1992). However, most of the extant diversity is relatively young (<60mya; Wood *et al.*, 2020) and a product of recent diversification where hybridization and polyploidization are thought to play an important role in species generation (Taylor & Hickey, 1992; Hoot *et al.*, 2004; Dai *et al.*, 2020). *Isoëtes* is an ideal study system for investigating the evolutionary implications of hybridization and polyploidization within vascular plants because of its unique evolutionary dynamics (i.e., an old clade with recent diversification), high species richness, pervasiveness of polyploids, and complex biogeographical patterns.

Although there is great potential for investigating reticulate evolution within *Isoëtes*, discerning species relationships in the genus has historically proven difficult due to the conserved body plan with limited morphological characters and character states (Pfeiffer, 1922; Hickey *et al.*, 1989). In fact, early Isoetologist W. N. Clute (1905) lamented, *“the marks by which the species [of* Isoëtes*] are distinguished are so obscure as to be puzzling to all but the select few, and in consequence the species have been largely taken upon by faith.”* In addition, recent work demonstrates that low molecular divergence further compounds the difficulties of circumscribing species within this genus (Pereira *et al.*, 2017; Wood *et al.*, 2020).

One species complex of *Isoëtes* that has remained particularly challenging is known from western North America. This complex includes two diploid progenitors (*I. echinospora* Durieu and *I. bolanderi* Engelm.), a sterile diploid hybrid (*I.* ✕ *herb-wagneri* W. C. Taylor), a fertile allotetraploid (*I. maritima* Underwood), fertile hexaploid (*I. occidentalis* L. F. Hend.), and a handful of sterile interploid hybrids (*I.* ✕ *pseudotruncata* D. M. Britton & D. F. Brunt. and *I.* ✕ *truncata* Clute; **Fig. 1, 2a**). The evolutionary relationships among taxa in this complex have been suggested before using morphology and cytology, but a consensus has not been reached (Fig. **1**; Britton & Brunton, 1993, 1996; Britton *et al.*, 1999; Taylor, 2002).

**Figure 1.**
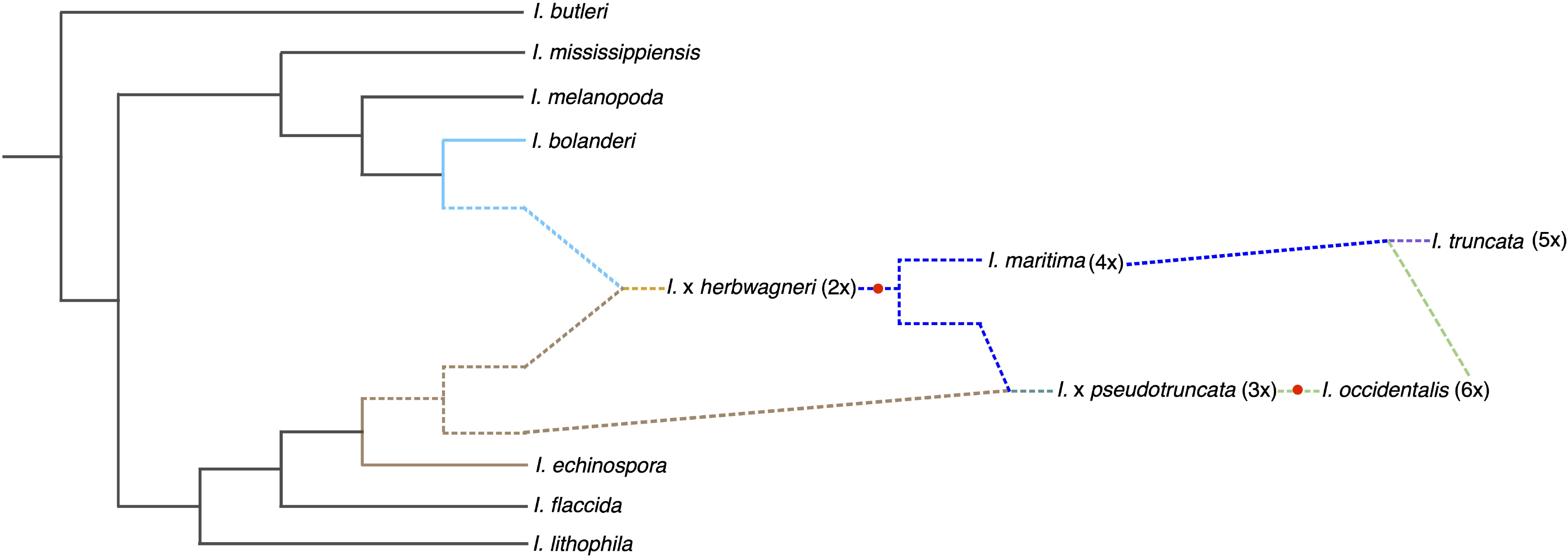
Historically hypothesized relationships of western North American *Isoëtes*, derived from Britton & Brunton, 1993, 1996; Britton *et al.*, 1999; and Taylor, 2002. *Isoëtes bolanderi* and *I. echinospora* hybridize to form the sterile diploid *I.* ✕ *herb-wagneri*. *Isoëtes maritima* is formed via a whole genome duplication (WGD; indicated by red dot) of *I.* ✕ *herb-wagneri.* Then, *I. maritima* (2x gamete) hybridizes with *I. echinospora* (1x gamete) to form triploid *I.* ✕ *pseudotruncata*. The triploid goes through a WGD (red dot) to form *I. occidentalis*. Finally, *I. maritima* (2x gamete) and *I. occidentalis* (3x gamete) hybridize to form the pentaploid *I. truncata*.

**Figure 2.**
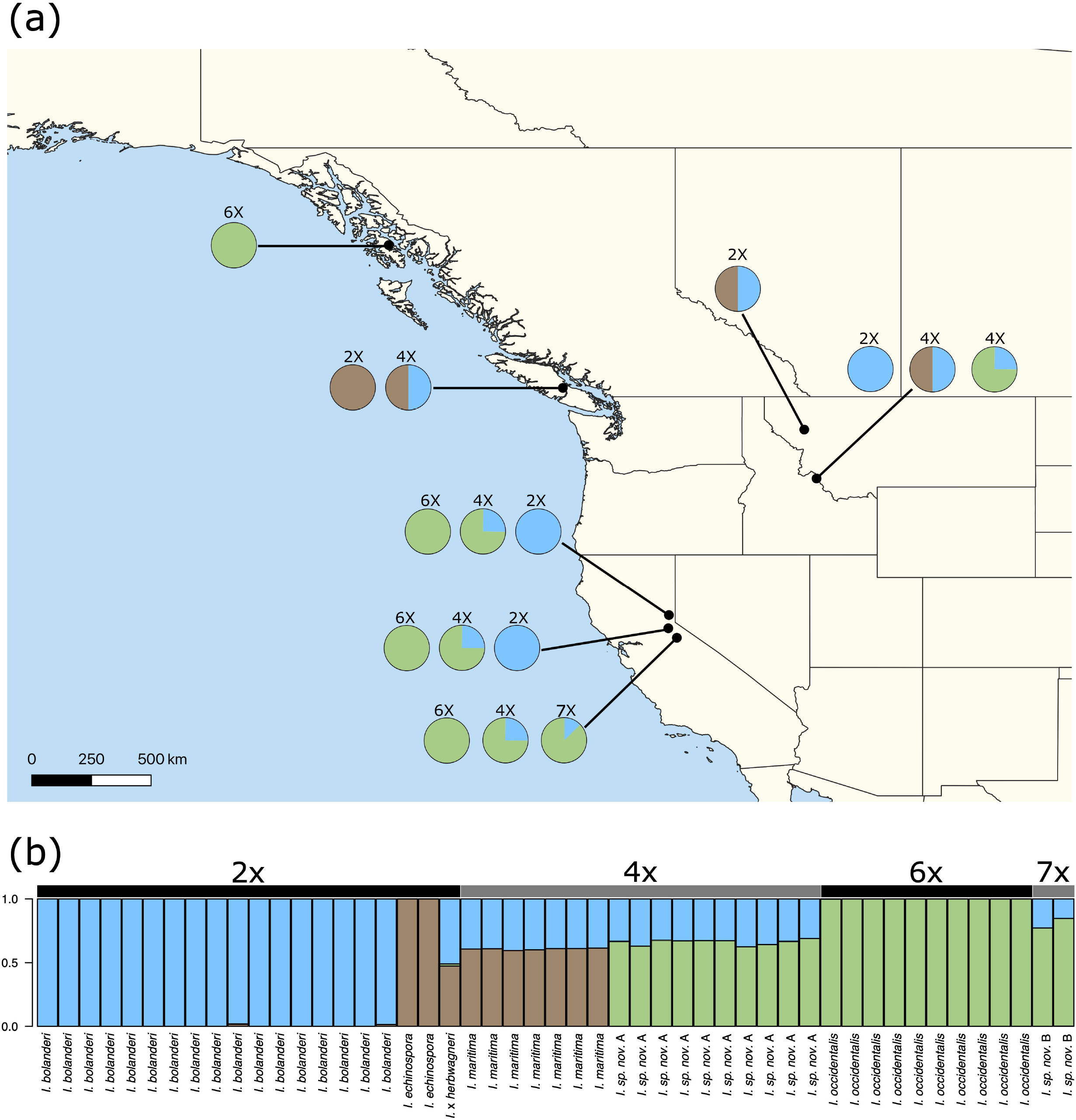
Sampling locations and population structure (*K* = 3) of western North American *Isoëtes*; (a) black dots represent lake localities where species were collected, pie charts indicate the ploidy and genomic structure of the different individuals collected at each location. The ploidy was derived from c-values or chromosome counts and the genomic constitution derived from the STRUCTURE analysis; (b) population structure and ploidy level for each sample; the diploid species (*I. bolanderi*, blue; *I. echinospora*, brown) and hexaploid I. occidentalis (*green*) are separated as the three main species with distinct genomic populations.

Here, we revisit this complex with new data and use phylogenomic comparisons to investigate the evolutionary implications of polyploidy and hybridization. We leverage five distinct data sets to accomplish this: low copy nuclear markers, whole chloroplast genomes, restriction site-associated DNA sequencing (RADseq), cytology (genome size estimates and chromosome counts), and reproductive status (fertile or sterile) from spore morphology. With these data, we are able to reconstruct the reticulate evolutionary history and directionality of hybridization events, as well as ploidy and fertility of hybrid taxa. This is the first phylogenomic study investigating the evolutionary history of a polyploid complex in *Isoëtes.* These data provide robust evidence that hybridization and polyploidy occur with frequency in the genus and that allopolyploidy—and potentially autopolyploidy—may be important evolutionary processes shaping extant *Isoëtes* diversity.

## Materials and Methods

We collected five datasets to investigate the reticulate relationships in a complex of western North American *Isoëtes*. We performed flow cytometry to estimate genome size; utilized Illumina short-read ddRADseq to obtain genomic data for population structure analysis and phylogenetic network analysis; identified fertile vs. aborted spores; sequenced whole chloroplasts with Illumina; and sequenced all alleles of the *LFY* nuclear marker with the Pacific Biosciences long-read platform to perform phylogenetic analyses.

### Taxon sampling

We assembled five distinct datasets, with collections from various sources. For RADseq and cytology, we collected fresh and silica dried leaf tissue from localities with historical information for each taxon across western North America (Fig. 2a). These included high-elevation lakes in Montana, California, and British Columbia. To determine specimen fertility, we examined spores from these collections. Specimen voucher information can be found in (Supplementary Table **S1**). For chloroplast genome sequencing and *LFY* allele sequencing we sampled from silica dried specimens of vouchered specimens (Table **S2**).

### Cytology

Fresh leaf material was used to measure C-values for individuals of each taxon in order to estimate their ploidy. C-values were estimated using microphylls from living plants. Leaves were prepared for DNA flow cytometry as described in Bolin *et al.*, (2017) using propidium iodide stain (P121493; Molecular Probes FluoroPure, ThermoFisher Scientific, Waltham, Massachusetts, USA). Newly expanded leaves of the *Glycine max* ‘Polanka’ (Doležel *et al.*, 1994) were used as the standard for genome size estimation. Samples were analyzed with a BD Accuri C6 Flow Cytometer and associated software (Becton Dickinson and Company, Franklin Lakes, New Jersey, USA), using gating and peak estimation parameters described in Bolin *et al.*, (2017). C-values were calculated for each individual following the methods of Dolezel & Bartos, (2005). By measuring the C-value for taxa with well documented chromosome counts we were able to infer ploidy levels for individual plants using C-value estimates.

### RADseq DNA extraction, library construction, and sequencing

Silica-dried plant tissue samples were supplied to the University of Wisconsin-Madison Biotechnology Center for DNA extractions, library preparation, and sequencing. DNA was extracted using the QIAGEN DNeasy mericon 96 QIAcube HT Kit and quantified using the Quant-iT™ PicoGreen dsDNA kit (Life Technologies, Grand Island, NY).

Libraries were prepared following Elshire *et al.*, (2011) with minimal modification. 100 ng of DNA were digested using PstI and MspI (New England Biolabs, Ipswich, MA) after which barcoded adapters amenable to Illumina sequencing were added by ligation with T4 ligase (New England Biolabs, Ipswich, MA). The 96 adapter-ligated samples were pooled and amplified to provide library quantities amenable for sequencing, and adapter dimers were removed by SPRI bead purification. Quality and quantity of the finished libraries were assessed using the Agilent Bioanalyzer High Sensitivity Chip (Agilent Technologies, Inc., Santa Clara, CA) and Qubit dsDNA HS Assay Kit (Life Technologies, Grand Island, NY), respectively. Size selection was performed to obtain 300 - 450 BP fragments. Sequencing was done on Illumina NovaSeq 6000 2×150 S2. Data were then analyzed using the standard Illumina Pipeline, version 1.8.2.

### RADseq data processing

All RADseq data processing was done on the Pearse Lab (Utah State University) high-performance workstation; downstream analyses were performed on the University of Utah Center for Higher Performance Computing. Raw data were demultiplexed using *stacks* v. 2.4 *process_radtags* (Catchen *et al.*, 2011, 2013) allowing for a maximum of one mismatch per barcode. Demultiplexed FASTQ files were paired and merged using *ipyrad* v. 0.9.52 (Eaton & Overcast, 2020). Low quality bases, adapters, and primers were removed from each read. Filtered reads were clustered at 90% similarity, and we required a sequencing depth of 6 or greater per base and a minimum of 35 samples per locus to be included in the final assembly. The *ipyrad* pipeline defines a locus as a short sequence present across samples. From each locus retained in the final assembly, *ipyrad* identifies single nucleotide polymorphisms (SNPs); these SNPs are the variation used in our downstream analyses. Although many of the species of *Isoëtes* included in this study are polyploid, *stacks* and *ipyrad* assume diploidy, so we processed all samples as such.

### Population structure analysis

The program STRUCTURE v. 2.3.4 (Pritchard *et al.*, 2000) estimates admixture (hybridization) between populations of closely related taxa. The program operates with the assumption that each individual’s genome is a mosaic from *K* source or ancestral populations. In our analysis, we ran STRUCTURE for *K* = 2 - 5 with 50 chains for each *K*. We then used CLUMPAK (Kopelman *et al.*, 2015) to process the STRUCTURE output and estimate the best *K* values (Pritchard *et al.*, 2000; Evanno *et al.*, 2005).

### Split network analysis

Split networks are implicit representations of reticulate evolution. They incorporate uncertainty by depicting multiple phylogenetic hypotheses: internal nodes do not represent ancestral taxa, but rather incongruencies between potential evolutionary histories. We utilized the NeighborNet (Bryant & Moulton, 2002) split network algorithm in SplitsTree v. 4.16.1 (Huson & Bryant, 2006). NeighborNet is similar to a neighbor joining tree in that it pairs samples based on similarity, sorting taxa into larger and larger groups (Bryant & Moulton, 2002). NeighborNet creates a network where parallel lines indicate splits of taxa, and boxes created by these lines indicate conflicting signals (Bryant & Moulton, 2002).

### Identification of fertility

Spores from sequenced individuals were photographed under a Zeiss Discovery v12 instrument (Carl Zeiss Microscopy, LLC, Thornwood, NY, USA). Some individuals were too immature to have viable spores and were not included. Individual spores were visualized so as to detect any abnormalities that would indicate sterility, such as flattening of proximal hemisphere, polymorphism, irregularity in size and texture, and connections of meiotic tetrads (Wagner *et al.*, 1986).

### DNA extraction for Illumina and PacBio sequencing

For whole plastome and nuclear gene sequencing, total genomic DNA was isolated from silica dried leaf tissue using the DNeasy Plant Mini Kit (Qiagen Inc., Valencia, CA, USA) at the Smithsonian NMNH Laboratory of Analytical Biology (LAB). DNA quantity and quality were measured using a Qubit 2.0 fluorometer (ThermoFisher Scientific, Waltham, MA, USA) and an Epoch spectrophotometer (BioTek Instruments Inc., Winooski, VT, USA).

### Chloroplast library preparation & sequencing

Whole genomic DNA was quantified using a Qubit 2.0 fluorometer (ThermoFisher Scientific, Waltham, MA, USA) and quality was measured by 260nm: 280nm absorption ratio using an Epoch spectrophotometer (BioTek Instruments Inc., Winooski, VT, USA). 150-1000ng of whole genomic DNA was diluted to 60μL and sheared to ~500BP fragments using the Q800R2 sonicator (Qsonica LLC, Newtown, CT, USA). Fragmented DNA was then prepared for sequencing on a MiSeq Illumina sequencer (Illumina Inc., San Diego, CA, USA) using the NEBNext Ultra II DNA Library Prep Kit for Illumina (New England Biolabs Inc., Ipswich, MA, USA). Manufacturers’ instructions were followed for end repair, adaptor ligation, indexing, and PCR enrichment for each sample. Libraries were quantified using a ViiA™ 7 Real-Time PCR System (Applied Biosystems Corp., Foster City, CA, USA). Libraries were then pooled and diluted to 4 nM and submitted for sequencing.

### Chloroplast genome assembly and phylogenetic inference

Illumina’s BaseSpace database was used to download index separated raw reads. Cutadapt (Martin, 2011) was used to trim NEB indices and filter out low quality bases. Pairing, assembling and referencing the chloroplast reads were done according to Schafran *et al.*, (2018). Adapters and low quality bases were trimmed using Trimmomatic 0.39 (Bolger et al. 2004). Paired reads were then mapped to the reference plastome of *Isoëtes flaccida (Karol et al., 2010)* using Bowtie2 (Langmead & Salzberg, 2012), to extract chloroplast reads. The putative chloroplast reads were then *de novo* assembled using SPAdes 3.10.1 with k-mer lengths 21, 33, 55, 66, 99, and 127bp (Bankevich *et al.*, 2012). Reference-based assemblies were constructed in Geneious Prime 2020.2.3 (https://www.geneious.com) by mapping putative chloroplast reads to the *I. flaccida* plastome and extracting the majority consensus sequence. For each sample, *de novo* assembled contigs were aligned to the reference-based assembly and any discrepancies were manually corrected using mapped reads. Whole plastomes were aligned with previously sequenced plastomes (Schafran *et al.*, 2018) using MAFFT in Geneious (Kearse *et al.*, 2012) and trees were generated using RAxML 7.3.0 (Stamatakis, 2006) and MrBayes 3.2.6 (Ronquist & Huelsenbeck, 2003). The GTR+G model, with 4000 bootstrap iterations was implemented in RAxML. All MrBayes analyses were run for 50 million generations, sampling every 1,000 generations, with one cold chain and three heated chains. MCMC output was visualized in Tracer 1.7 to check for convergence among the chains (Rambaut *et al.*, 2018). These computations were run on the FASRC Odyssey cluster supported by the FAS Division of Science Research Computing Group at Harvard University.

### PacBio library preparation

In preparation for PacBio sequencing the second intron of the nuclear gene *LEAFY* (*LFY*) was amplified by the polymerase chain reaction (PCR), using modified 30F (5’-GATCTTTATGAACAATGTGG-3’) and 1190R (5’GAAATACCTGATTTGTAACC3’) primers. Amplifications in 25μL reactions were conducted using the following reagents: 12.5μL GoTaq (Taq polymerase master mix), 0.5μL of 0.1mM BSA (Bovine Serum Albumin), 1μL of DMSO (Dimethyl Sulfoxide), 1μL of 10μM Forward and Reverse primer, respectively, 6.5μL of Nuclease free H_2_O, and 2.5μL of DNA. PCR protocol began with an initial denaturation temperature of 94°C for 4 minutes, 10 cycles of 94°C for 1 minute, 58°C for 1 minute and 72°C for 1 min, followed by 25 cycles of 94°C for 30 seconds, 53°C for 1 minute, and 72°C for 30 seconds. A final extension cycle of 72°C for 4 minutes followed the reaction. *LFY* amplicons were bead cleaned using Kapa beads following the (manufacturer’s protocol). A 1:1 ratio of bead solution to amplicon was used to size select for ~1100bp fragments. The solution was resuspended in 20μL of nuclease free H_2_O. Each cleaned sample was then proportionally pooled, based on concentration and ploidy-level into a 1.5ml microcentrifuge tube and sent to the Duke University Sequencing and Genomic Technologies Shared Resource (Durham, NC) for PacBio RSII sequencing using two SMRT cells.

### PacBio data processing and phylogenetic inference

Circular consensus sequences (CCS) provided by the Duke University Sequencing and Genomic Technologies Shared Resource were size filtered to the expected size of the amplicon (1000-1200 bp) and then demultiplexed and clustered with PURC (Rothfels *et al.*, 2017; Dauphin *et al.*, 2018). Demultiplexed reads were also filtered by DADA2 (Callahan *et al.*, 2016) to remove those with any “N” base calls and with more than 5 expected errors. Amplicon sequence variants (ASVs) were identified from the filtered, demultiplexed reads also using DADA2. Mock community data show that DADA2 more reliably recovers the true composition of a mixture of amplicon CCS reads compared to OTU clustering, which tends to oversplit clusters (Nelson *et al.*, 2020a). Therefore, we use only ASVs in downstream analyses. ASVs were aligned to a selection of published *Isoëtes LFY* sequences (Hoot *et al.*, 2004; Taylor *et al.*, 2004; Kim *et al.*, 2010; Rosenthal & Rosenthal, 2014) using the G-INS-i algorithm in MAFFT V7.313 (Katoh & Standley, 2013) and phylogenies were inferred using MrBayes (Ronquist & Huelsenbeck, 2003) using the parameters described above for chloroplast phylogenetic inference.

## Results

We initially identified all specimens based on morphology (Pfeiffer, 1922; Britton & Brunton, 1993, 1996; Taylor, 2002), although some specimens could not be assigned to a named species. Our morphological identifications were compared to our molecular results (plastid, LFY nuclear marker, and RADsq data; discussed in detail below), and updated accordingly. Most specimens were ultimately identified as named taxa. However, we detected two taxa (with a combination of cytology, molecular data, and examination of spores) that did not match any named species. These are hereafter referred to as *Isoëtes species novo* A (tetraploid) and B (septaploid). Further morphological data are needed for formal taxonomic description.

### Cytology

C-values were only obtained from specimens collected with fresh leaf tissue, for this reason genome size was not measured for every individual in this study (Table **S1**). Ploidy was inferred from C-values based on the relationship between chromosome number and C-value (Bolin *et al.*, 2017). For other individuals, chromosome counts have been made. As expected, C-values of the putative diploids, *I. echinospora* and *I. bolanderi* were the smallest. The values of all samples of *I. bolanderi* were between 2.4–2.6, while the value for *I. echinospora* was 3.9 (Table **S1**). Specimens of *I. echinospora* from eastern North America are also known to have a larger C-value compared to other *Isoëtes* diploids, and the genome size of diploid *Isoëtes* species can vary (Bolin *et al.*, 2017). The values of the tetraploids *I. maritima* as well as *I. sp. nov.* A are between 5.7–6.9, and the values of the hybrid between *I. occidentalis* and *I. bolanderi* are between 5.7–6.9 (Table **S1**). C-values of the hexaploid *I. occidentalis* ranged from 9.4–9.8, and values for *I. sp nov.* B (7x) had a value between 9.9–10.8 (Table **S1**). Generally, there is a tight correlation between C-value and ploidy, but within *Isoëtes* this relationship starts to break down at higher ploidy levels (Bolin *et al.*, 2017). Because our lower ploidy level taxa have relatively high C-values, the high values of the tetraploids, hexaploids, and septaploids are mathematically consistent. For instance, as *I. maritima* (average C-value: 6.2) is a hybrid between *I. echinospora* (3.9) and *I. bolanderi* (average C-value: 2.5), simple addition of the genomic size of these two taxa corroborates the larger genome size found in our polyploids.

### Population genomic analysis

Our raw data obtained from the University of Wisconsin-Madison Biotechnology Center consisted of an average of 2.33 × 10^6^ raw reads per sample. In the final assembly, we retained an average of 2,980 loci per sample, with a standard deviation of 858 loci per sample. Our STRUCTURE and SplitsTree analyses utilized a dataset of 3,343 SNPs, composed of one SNP per locus in the final assembly. Raw, demultiplexed sequences can be accessed from the NCBI GenBank Sequence Read Archive (PRJNA665237).

To identify the optimal number of populations (*K*) in our population genomic analysis, we used the best *K* method (Evanno *et al.*, 2005), as well as visually examining *K* = 2-5 as recommended by Pritchard *et al.* (2000). The best *K* method indicated a value of 3 as the best fit for our data (Fig. **S1**), which agreed with our prior morphological and cytological hypotheses on the genomic makeup of these taxa (Fig. **2b**). This was also the most biologically meaningful number of groups because it separated each named diploid, and elucidated parentage in the allopolyploid hybrids. Decreasing *K* below 3 leads to a loss in information and increasing it only adds noise and does not add any meaningful population clusters (Fig. **S1**). Diploids identified as *I. bolanderi* and *I. echinospora* fall into respective single population groups (Fig. **2b**). Putative hybrids including *I.* ✕ *herb-wagneri* and *I. maritima* cluster together with half of their genomic constitution coming from *I. echinospora* and *I. bolanderi*. *Isoëtes* ✕ *herb-wagneri* is a sterile diploid hybrid, while *I. maritima* is a fertile allotetraploid. Based on population structure analyses, the genomic constitution of *I. maritima* has slightly more *I. echinospora* ancestry than *I. bolanderi* (Fig. **2b**), but this could be accounted for by sequencing or analysis errors in the polyploid taxon.

All samples of the fertile hexaploid *I. occidentalis* clustered together as a distinct population, without signatures of any other species (Fig. **2b**). We recovered a sterile hybrid cross between *I. occidentalis* and *I. bolanderi* (*I. sp. nov.* A). Based on cytology this taxon is likely to be a tetraploid. There was an additional septaploid hybrid between *I. occidentalis* and *I. bolanderi* (*I. sp. nov.* B) (Fig. **2b**).

Our split network analysis showed very similar groups as our population structure analysis. *Isoëtes echinospora* and *I. occidentalis* fall into two separate, but closely-related groups; *Isoëtes bolanderi* is also placed as a distinct group, but farther away from the latter two species (Fig. **S2**). *Isoëtes* ✕ *herb-wagneri* and *I. maritima* sit very close to one another, further substantiating *I. maritima* as an autopolyploid derivative of *I.* ✕ *herb-wagneri*. These two species are placed in between their hypothesized progenitors, and the boxes created by the split network indicate some conflicting evolutionary signals (i.e., hybridization). The allopolyploid taxa *I. sp. nov.* A is placed in between *I. bolanderi* and *I. occidentalis*. *Isoëtes sp. nov.* B appears both next to the former hybrid, and with *I. occidentalis*. Due to its similarity to the population structure analysis (Fig. **S2**).

### Determination of spore viability

To determine whether a taxon is fertile or sterile, and thus whether it is of hybrid origin, megaspores were visualized. For each mature individual, megaspores were visualized under a dissecting scope. We followed Wagner *et al*. (1986) to identify whether spores are spherical, uniform in size and ornamentation indicating viability and fertility, or flattened, uneven in size and ornamentation indicating non-viable and sterility (Wagner *et al.*, 1986). We determined that most of the spores from *I. sp nov A., I. sp. nov. B.*, and *I.* ✕ *herb-wagneri* were variable in shape and size, suggesting sterility, while spores from the diploid species (*I. echinospora* and *I. bolanderi*), the tetraploid *I. maritima*, and the hexaploid *I. occidentalis* were uniform in size and shape, suggesting fertility (Fig. **3a-g**).

**Figure 3.**
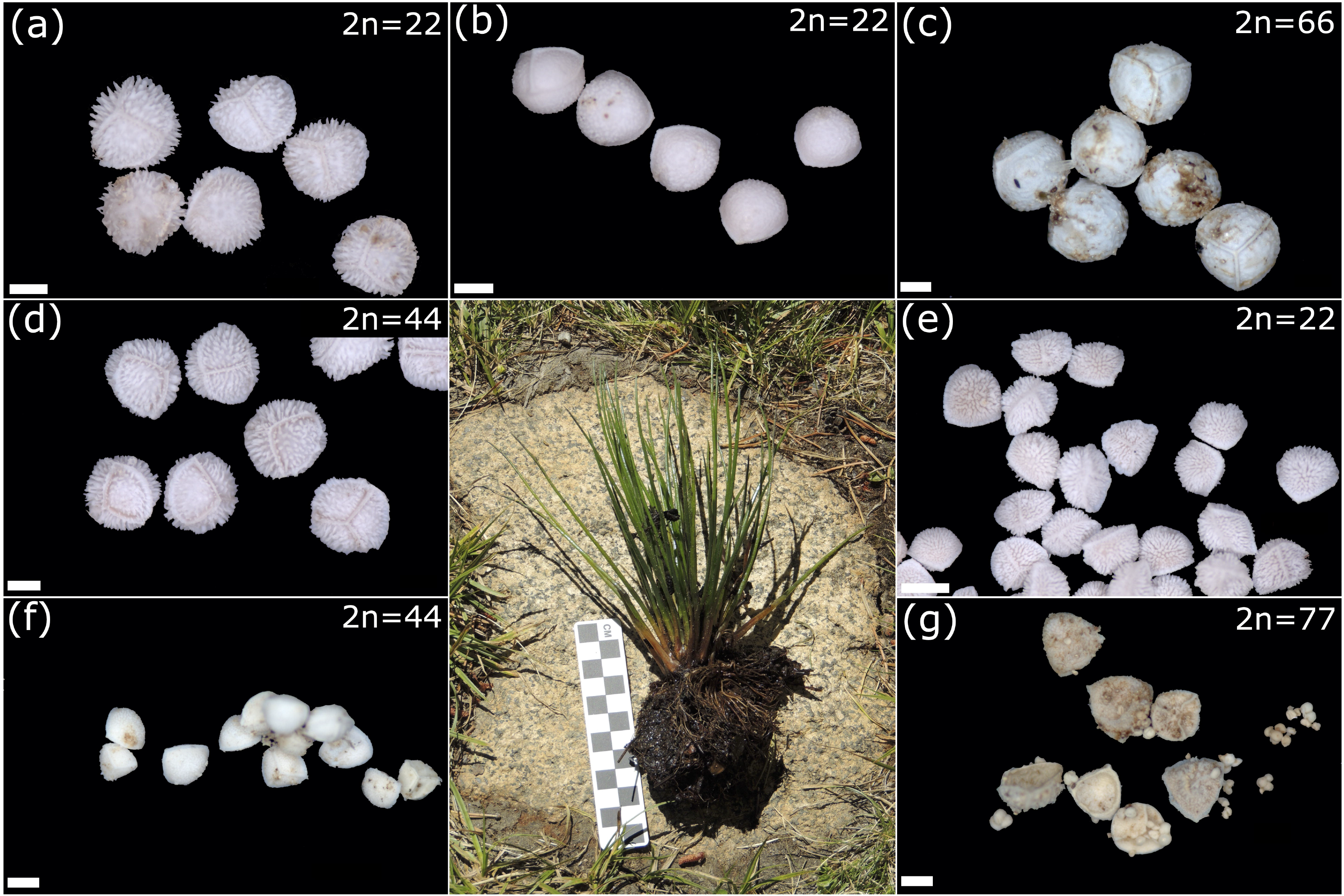
Spore images for (a) fertile diploid *Isoëtes echinospora*; (b) fertile diploid *I. bolanderi*; (c) fertile hexaploid *I. occidentalis*; (d) fertile tetraploid *I. maritima*; (e) sterile diploid *I.* ✕ *herb-wagneri*; (f) sterile tetraploid *I. sp. nov.* A; and (g) sterile septaploid *I. sp. nov.* B. (scale bar: 200μm). Fertile spores were round and uniform within the sample (i.e., a, b, c, d). Sterile spores are misshapen (sometimes flattened: e) and vary in size sometimes with the meiotic tetrads visible (i.e., f, g). Central image is of *I. occidentalis* in the field.

### Whole chloroplast genomes

In order to gauge the directionality of hybridization events whole plastome phylogenies were reconstructed (Fig. **4**). Bootstrap support as well as posterior probability were high (>95; >0.95) for almost every node in the plastome-derived phylogeny (Fig. **4**). Three primary clades of interest were recovered: an *I. bolanderi* clade (blue); an *I. echinospora* clade (brown); and an *I. occidentalis* clade (green). The branch leading to each of these clades has a posterior probability and bootstrap support of 1.00 and 100% respectively. The *I. echinospora* and *I. occidentalis* clades fall out as sister groups to each other, while the *I. bolanderi* clade is sister group to *I. melanopoda* (Fig. **4**). Two individuals of *I. maritima* appear in different places in the tree with one as sister to *I. echinospora*, while the other appears most closely related to *I. bolanderi*, suggesting that each diploid species can be the maternal parent. Individuals identified in the field as *I.* ✕ *pseudotruncata*, as well as the hybrid *I. sp. nov.* A are most closely related to *I. occidentalis*. No diploid progenitor was found in the *I. occidentalis* clade. Similar to *I. maritima* two individuals of *I.* ✕ *herb-wagneri* appear most closely related to either *I. bolanderi*, or *I. echinospora*, suggesting that either parent can be the maternal progenitor in those hybridization events (Fig. **4**).

**Figure 4.**
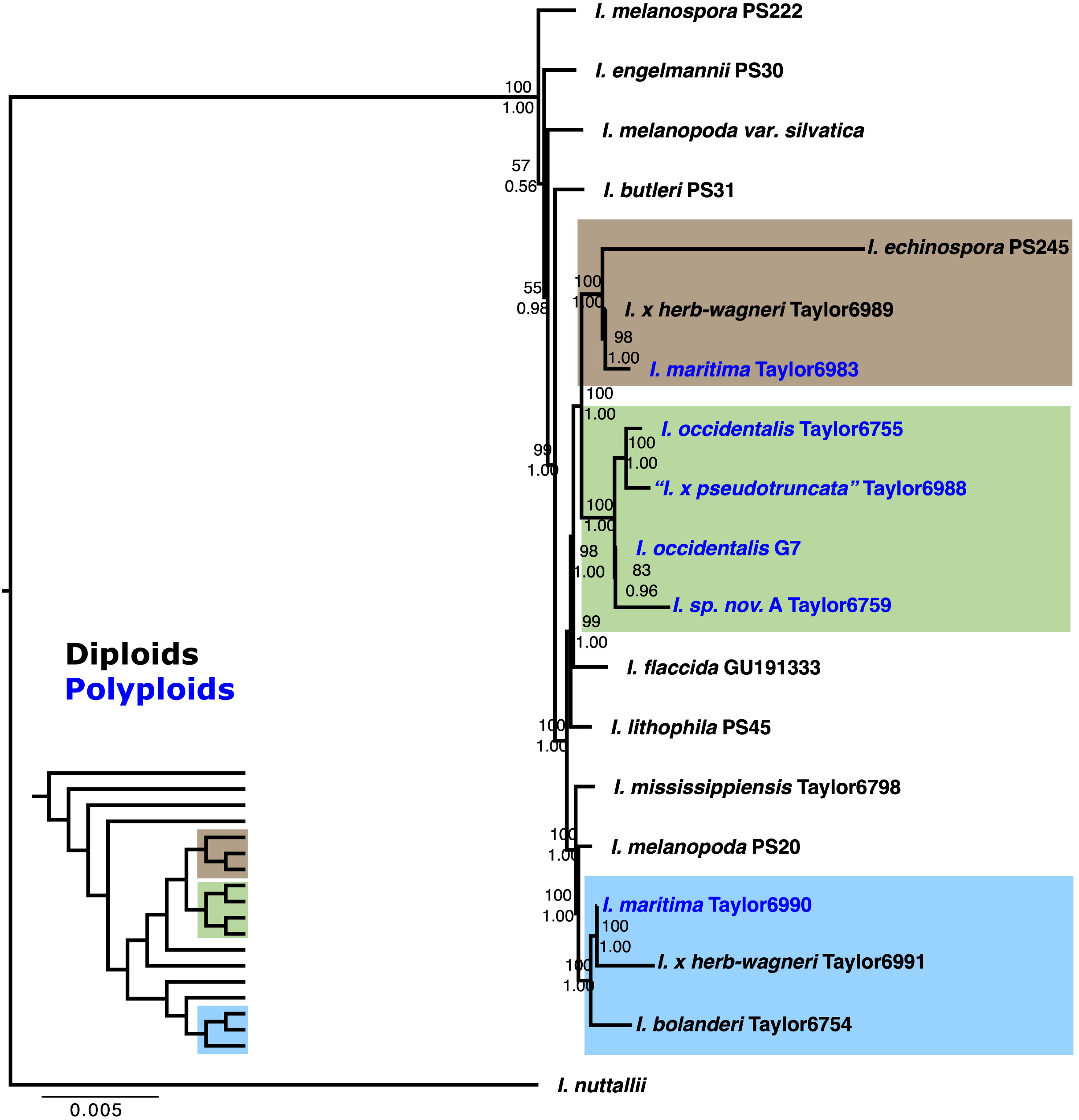
Plastome phylogeny of western North American *Isoëtes* including outgroups for reference. All individuals in the group of interest comprise three distinct clades: *I. bolanderi* (blue), *I. echinospora* (brown), *I. occidentalis* (green). Within the blue clade, there are accessions of *I. bolanderi*, *I.* ✕ *herb-wagneri*, and *I. maritima*. In the green clade, there are accessions of *I. occidentalis* and *I. sp. nov.* A. Finally, in the brown clade there are accessions of *I. echinospora*, *I.* ✕ *herb-wagneri*, and *I. maritima*. The taxon labeled “*I. pseduotruncata*” Taylor 6988, was initially hypothesized to be *I. pseduotruncata*, however our data suggests this is *I. sp. nov* A.

### LFY phylogenetic inference

PacBio sequencing yielded 5684 CCS reads. Despite efforts to achieve equal sequencing depth, the number of reads per sample ranged from 10 to 926 (average 334, standard deviation 228). Following filtering, the number of reads per sample decreased by an average of 43%. DADA2 identified a total of 29 ASVs compared to the 39 OTU clusters created with PURC. Two examples of disagreement in the number of ASVs recovered from five pairs of PCR replicates suggests that PCR or sequencing bias may still affect results, although one of these disagreements may be explained by very low coverage of one replicate (number of filtered reads = 3).

These single gene nuclear data provided the initial framework we used to examine this reticulate complex. Each taxon sampled using PacBio’s RSII platform returned a single sequence or multiple sequences depending on the ploidy level for each taxon (i.e. diploids returned single sequences while allopolyploids returned multiple). The PURC pipeline occasionally outputs multiple clusters of the same sequence due to strictness of clustering parameters (Fig. **S3**). The output from the PURC pipeline shows multiple clusters for *I.* ✕ *herb-wagneri*, *I. maritima*, multiple unnamed hybrids, and the two replicates of *I. occidentalis* (Fig. **S3**). The hypothesized hybrid (*I.* ✕ *herb-wagneri*) and allotetraploid (*I. maritima*) shows equal clustering of *LFY* alleles from both putative parent species (*I. bolanderi* and *I. echinospora*). The multiple unnamed taxa (denoted as ‘sp.’) are not yet determined taxonomically and will be dealt with in a later paper. Various individuals of *I. echinospora* are found in multiple places in the phylogeny, indicating either misidentification or analytical errors (Fig. **S3**). The node splitting *I. occidentalis* and *I. echinospora* has very low support (0.56) and most of the tree is not resolved (i.e., soft polytomy) (Fig. **S3**). However, this is expected when using a single gene to reconstruct a species tree with extremely closely related and reticulating taxa.

## Discussion

Reticulate evolution is thought to be common in the genus *Isoëtes* because of the frequent occurrence of sterile plants, various co-occurring cytotypes, and the high proportion of polyploid taxa (Taylor & Hickey, 1992; Troia *et al.*, 2016). Previous attempts to explain the patterns of hybridization and polyploidization have relied on few genetic markers (Hoot *et al.*, 2004; Kim *et al.*, 2010; Pereira *et al.*, 2018; Dai *et al.*, 2020), which do not accurately depict species relationships in recently diverged groups with low molecular divergence, introgression, and incomplete lineage sorting (Nelson *et al.*, 2020b). Using whole chloroplast and nuclear genomes, in conjunction with cytological and reproductive data we investigated the importance of hybridization and polyploidization within a complex of western North American *Isoëtes*. We detected several hybrids and polyploids, their parental origin, and the directionality of hybridization events. Specifically, we find evidence for homoploid hybridization leading to the formation of sterile diploid taxa that undergo whole genome duplication resulting in fertile polyploid lineages (Fig. **2**, **5, S2, S3**). These allopolyploids can hybridize with their diploid progenitors, across ploidy levels, to form sterile interploid hybrids (Fig. **3**, **5**). Interestingly, the directionality of hybridization events is reciprocal between taxa of the same cytotype but in interploid hybrids, the higher ploidy taxon acts solely as the maternal donor (Fig. **4**). Finally, we find evidence for a distinct hexaploid lineage without signatures of hybrid origin, raising questions about its origin (Fig. **2b**, **4, S2**).

**Figure 5.**
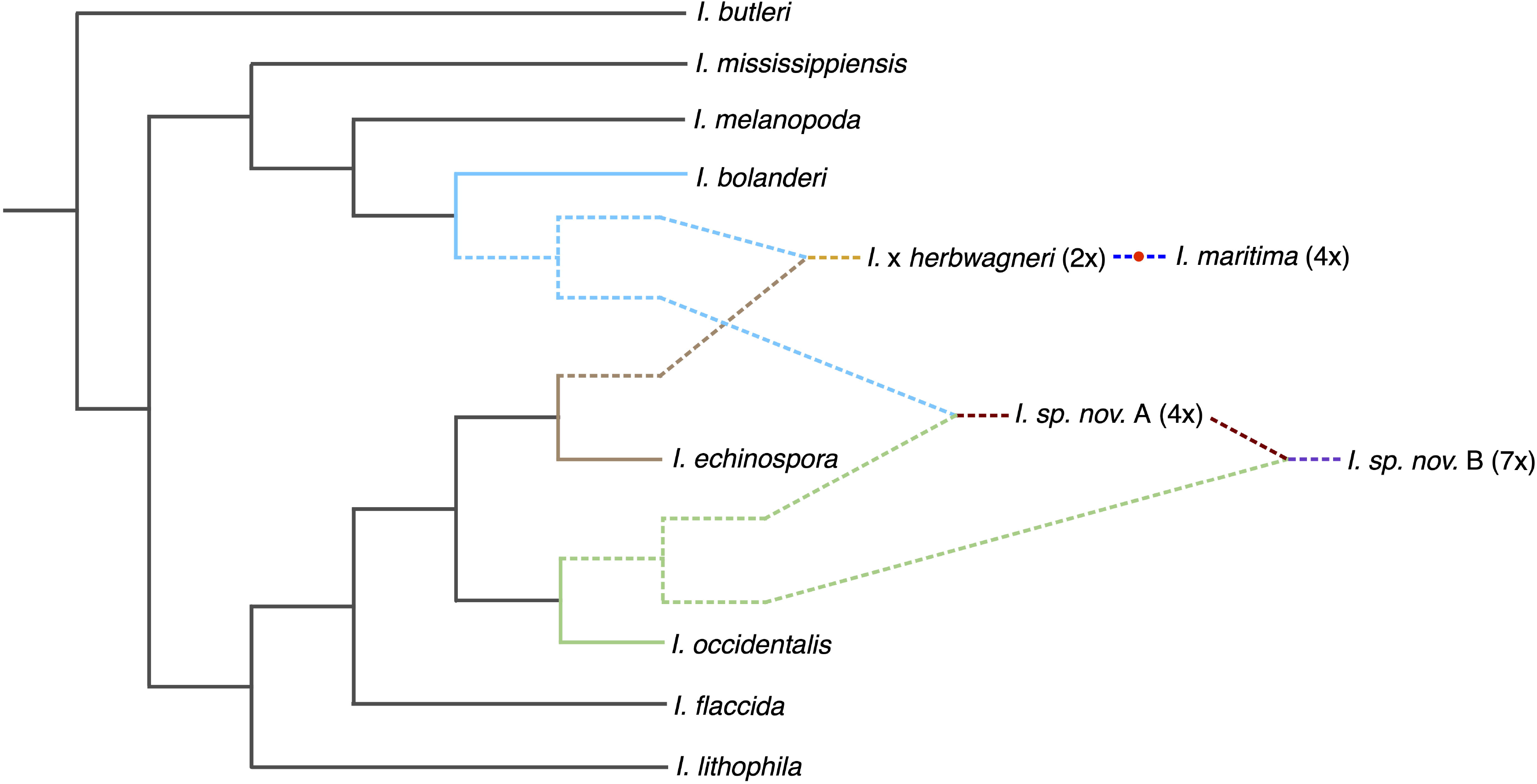
Hypothesized relationships of western North American *Isoëtes* as supported by this study. *Isoëtes bolanderi* and *I. echinospora* hybridize to form the sterile diploid *I.* ✕ *herb-wagneri*. *Isoëtes maritima* is formed via a whole genome duplication (indicated by red dot) of *I.* ✕ *herb-wagneri. Isoëtes bolanderi* (1x gamete) and *I. occidentalis* (3x gamete) hybridize to form the tetraploid *I. sp. nov.* A. Finally, *I. occidentalis* (3x gamete) and *I. sp. nov.* A. (4x gamete) hybridize to form the septaploid *I. sp. nov.* B. The polyploid origin of *I. occidentalis* is still inconclusive.

### Homoploid hybridization and allopolyploid speciation

The diploid species *Isoëtes bolanderi* and *I. echinospora* co-occur throughout western North America. We find that genomic constitution (Fig. **2b**, **S2, S3**), ploidy (Table **S1**), sterility (Fig. **3e**), and chloroplast inheritance (Fig. **4**), demonstrate that these species hybridize to form the sterile homoploid *I.* ✕ *herb-wagneri* (Fig. **5**), initially hypothesized by Taylor (2002). Since *I.* ✕ *herb-wagneri* is sterile (Fig. **3e**) it cannot reproduce sexually and, in the absence of asexual reproduction through rootstock division, each individual is a product of a new hybridization event. This means that *I.* ✕ *herb-wagneri* cannot persist as a sexually reproducing independent lineage, a necessary step in homoploid hybrid speciation (Yakimowski & Rieseberg, 2014). Based on these data, it is clear that *Isoëtes bolanderi* and *I. echinospora* are too evolutionary divergent to form fertile homoploid hybrids, and allopolyploidy must be a necessary step to restore fertility (Buggs *et al.*, 2009).

Based on overlapping distributions and similar morphology, *I. maritima* was historically circumscribed as a variety of *I. echinospora* (Löve, 1962). Cytological differences later led to the segregation of *I. maritima* as a distinct tetraploid species (Brunton & Britton, 1999). The formation of viable spores (Fig. **3**), tetraploidy (Table **S1**), chloroplast inheritance (Fig. **4**), and genomic constitution (Fig. **2b**, **S3**) demonstrate that *I. maritima* is the allotetraploid derivative of *I.* ✕ *herb-wagneri* (Fig. **5**). The formation of *I. maritima* may be achieved in multiple ways including: the fusion of unreduced gametes in *I.* ✕ *herb-wagneri*; the fusion of unreduced gametes from the diploid parents (*I. echinospora* and *I. bolanderi*); or through a triploid bridge (i.e., the fusion of a reduced gamete of one parent with an unreduced gamete of another, followed by a backcross of that triploid hybrid with a reduced gamete from one of the diploid parents) (Ramsey & Schemske, 2002; Husband, 2004). We seldom find abnormal spores in diploid species, but frequently find them in hybrids (Fig. **3**; JSS, WCT, and PWS, pers. obs), suggesting that the first scenario (two unreduced gametes of *I.* ✕ *herb-wagneri*) may be most likely. The presence of *I. maritima* and the sterility of *I. ✕ herb-wagneri* suggests that allopolyploid speciation is a more common mechanism of hybrid speciation, compared to homoploid hybrid speciation, within this complex.

### High-ploidy hybrids

In the 1990s, two sterile *Isoëtes* hybrids (*Isoëtes* ✕ *pseudotruncata* and *I.* ✕ *truncata*) were documented in northwestern North America (Britton & Brunton, 1993, 1996; Britton *et al.*, 1999). *Isoëtes* ✕ *pseudotruncata* (3x) was described as the hybrid of *I. maritima* ✕ *I. echinospora*; and *I.* ✕ *truncata* (5x) was describe as the hybrid of *I. maritima* ✕ *I. occidentalis* (Fig. **1**). Our analyses do not recover any genomic signatures of these putative hybridization events described by Britton and Brunton (1993, 1996; **Fig. 1, 2b**). It is possible that we did not sample them. Alternatively, they may have gone extinct or their progenitors were misidentified. However, we did find signatures of two other hybrid taxa, a sterile tetraploid (*I. sp. nov.* A) and a sterile septaploid (*I. sp. nov.* B) (Fig. **2b**; Table **S1**). Both have genomic signatures of *I. bolanderi* and *I. occidentalis* (Fig. **2b**, **S2, S3**), and they have polymorphic spores, indicating hybrid sterility (Fig. **3f, g**). We suspect that the tetraploid *I. sp. nov.* A is a hybrid between a reduced gamete of *I. occidentalis* (3x) and a reduced gamete of *I. bolanderi* (1x) (Fig. **5**); while the septaploid *I. sp. nov.* B is either the backcross between an unreduced gamete of *I. sp. nov.* A (4x) and *I. occidentalis* (3x) (Fig. **5**), or the hybrid product between an unreduced gamete of *I. occidentalis* (6x) and a reduced gamete of *I. bolanderi* (1x).

Abnormal chromosomal behavior is common among interploid hybrids (Sigel, 2016). Chromosomal non-disjunction and uneven chromosomal segmentation can occur during meiosis leading to spores of various ploidy (Sybenga, 1996). Therefore, it is possible that these two interploid hybrids (*I. sp. nov.* A and *I. sp. nov.* B) could produce meiotic products with various cytotypes, resulting in pentaploid and triploid taxa like *I.* ✕ *truncata* and *I.* ✕ *pseudotruncata*, respectively. Given this phenomenon, it is possible that *I.* ✕ *truncata* and *I.*✕ *pseudotruncata* are not derived from their initially proposed hybridization events (Fig. **1**; Britton & Brunton, 1993, 1996), but rather represent multiple cytotypes derived from abnormal chromosomal behavior (Fig. **5**). Other lines of evidence may support this, specifically the various spore sizes in these taxa (some large, round and seemingly viable; Fig. **3f, g**), and observation that the chloroplast genome of a sterile triploid initially identified as “*I.* ✕ *pseudotruncata”* (Fig. **4**; Taylor 6988) is actually derived from *I. occidentalis* (Fig. **4**). It would be unlikely for this taxon to inherit its chloroplast from *I. occidentalis* if it were a hybrid between *I. maritima* and *I. echinospora*, as initially hypothesized (Fig. **1**; Britton & Brunton, 1993). However, more data including DNA sequences from type species are needed to conclusively determine the identity of *I.* ✕ *pseudotruncata* and *I.* ✕ *truncata*.

These interploid hybrids occur in mixed populations in roughly equal (or lower) densities than their progenitors (JSS, WCT, pers. obs) and also hybridize with them, yet still persist in the population. The minority cytotype exclusion theory postulates that less common cytotypes within a population will be outcompeted by the majority cytotype, unless they have different ecological strategies (Levin, 1975). It is possible that interploid *Isoëtes* fill a disparate niche from their progenitors, or perhaps they are so novel that such evolutionary dynamics are still unfolding, and in the future these hybrids may naturally go extinct.

### Directionality of hybridization events

Because *Isoëtes* is heterosporous, both micro- and mega-gametophytes must be present for fertilization to occur. Both male and female gametophytes, however, often co-occur in these populations (Fig. **2a**), so one of the main, but not exclusive (Schneller, 1981), prezygotic barriers is sperm size and archegonial neck size (Testo *et al.*, 2015). Chloroplast inheritance shows that different individuals of *I.* ✕ *herb-wagneri* and *I. maritima* are either more closely related to *I. bolanderi* or *I. echinospora*, meaning either parent can be the maternal donor (Fig. **4**). Since *I. echinospora* and *I. bolanderi* are diploids they should have similar sized organs (i.e., sperm, eggs, archegonia), making reciprocal hybridization possible (Haufler *et al.*, 1995; Sigel *et al.*, 2014). In contrast, the interploid hybrid *I. sp. nov.* A is derived from a hexaploid and a diploid (Fig. **2b**, **5**), with the hexaploid *I. occidentalis* as the sole maternal progenitor (Fig. **4**). This is most likely due to sperm size of the hexaploid being larger than the sperm of the diploid (Testo *et al.*, 2015) and may not fit down the narrower archegonial neck of the *I. bolanderi* gametophyte.

### The origin of Isoëtes occidentalis and patterns of biogeography

Individuals identified as *Isoëtes occidentalis* in this study comprise a widespread and genomically uniform taxon throughout western North America (Fig. **2a,b**). Using preliminary morphological and cytological data, we tested the hypothesis that *I. occidentalis* was an allohexaploid derived from *I. maritima* and *I. echinospora* (Fig. **1**). However, we do not find signatures of shared ancestry within the genome of *I. occidentalis* (Fig. **2b**). Based on these results, we suggest that *Isoëtes occidentalis* may be either an autopolyploid or allopolyploid derived from extinct or unsampled parents. In our population structure analysis at *K* = 2, we find that *I. occidentalis* has the same genomic constitution as *I. echinospora* (Fig. **S1**), and our chloroplast (Fig. **4**) and nuclear data (Fig. **S3**) suggest they are each other’s closest sampled relatives. This could mean that *I. occidentalis* is an ancient autopolyploid of *I. echinospora*; alternatively, *I. occidentalis* may be an ancient allopolyploid derived from *I. bolanderi* and *I. echinospora*, but enough time has elapsed to allow for genetic divergence and genomic rearrangement (Sigel, 2016), masking the genomic similarities between these taxa in the population structure analysis (Fig. **2b**). Future analyses including more individuals from geographically diverse populations and investigating ancestral genomic signatures of introgression should be conducted to more firmly determine the origin of *I. occidentalis* and other high-ploidy species in *Isoëtes*.

Given the distribution of *I. occidentalis* throughout the Sierra Nevada, Rocky Mountains, and Coast Range, glaciation likely played a role in its evolutionary and biogeographic history (Haufler *et al.*, 1995; Burnier *et al.*, 2009; Stein *et al.*, 2010; Sessa *et al.*, 2012a,b; Jorgensen & Barrington, 2020). Progenitors of *I. occidentalis* may have occurred throughout lakes in northwestern North America, and Quaternary glaciation cycles (Dyke & Prest, 1987; Hughes *et al.*, 1989; Mandryk *et al.*, 2001) could have led to extinction of the diploid parents, while the polyploid *I. occidentalis* survived in refugia like the ‘ice-free corridor’ in western Canada (Schweger, 1989; Dyke, 2005). If true, it is likely that *I. occidentalis* evaded extinction due to a wider distribution and fixed heterozygosity, which buffered populations from inbreeding depression during climatic fluctuations (Brochmann *et al.*, 2004).

Long-distance dispersal could be another aspect of gene flow and biogeographical patterns in these western North American *Isoëtes.* Unlike most homosporous ferns, which disperse their spores via air currents (Barrington, 1993), spore dispersal in *Isoëtes* is poorly understood because they are aquatic and produce large megaspores (200-1000μm), unable to be dispersed by wind. Anecdotal evidence suggests they can be dispersed by waterways, waterfowl, or even snails and earthworms (Duthie, 1929; Jermy, 1990; Taylor & Hickey, 1992; Larsén & Rydin, 2016), and recent work has documented spores in waterfowl fecal matter (Silva *et al.*, 2020), confirming previous observations. It is not inconceivable that sporangia, spores, or even whole plants are dispersed in the crops of these migratory animals (Les *et al.*, 2003). In this way, separate species that evolved in allopatry, with limited reproductive barriers, could be re-introduced by waterfowl, leading to hybridization. We suspect that long-distance dispersal through waterfowl and glaciation cycles have also influenced biogeographical patterns of other *Isoëtes* species, and further work should aim to tease apart these processes. Our results demonstrate that hybridization and polyploidization are common and occur with ease in this complex of western North American *Isoëtes*. The occurrence of many polyploids and putative hybrids in the genus (Troia *et al.*, 2016) suggests that these processes are important in shaping extant *Isoëtes* diversity.

## Acknowledgements

Thank you to William D. Pearse for thoughtful comments on the manuscript. We also thank Gabriel Johnson and undergraduate research assistance from Catawba College for help with laboratory procedures and C-value generation. We thank the University of Wisconsin Biotechnology Center DNA Sequencing Facility for providing DNA extraction, library prep, and DNA sequencing facilities and services. Parts of this project were funded by The Society of Systematic Biologists Graduate Student Research Award, awarded to JSS. SPK is funded by a National Science Foundation Graduate Research Fellowship.

## Author contributions

Taylor and Zimmer initiated the project, Suissa and Taylor carried out fieldwork, Bolin determined C-values, Suissa and Schafran sequenced and analyzed plastid genomes and *LFY* sequences, Suissa and Kinosian performed RADSeq data processing, Kinosian performed split network and population genomic analyses, and all authors participated in writing the manuscript.

## Data availability

All sequence data can be accessed from the NCBI Genbank Sequence read archive. Demultiplexed sequences from RADSeq: PRJNA665237.

