## Supplemental Information for "Revealing the evolutionary history of a reticulate polyploid complex in the genus *Isoëtes*"

***Phytologist* Supporting Information**

Article acceptance date: 30 October 2020

The following Supporting Information is available for this article:

**Fig. S1** Population structure analysis for western North American *Isoëtes* at *K* = 2-5. Unique specimen names are indicated at the bottom of the figure, species names and ploidy are at the top of the figure. *K* = 3 is the most biologically meaningful arrangement of populations. The diploid species (*I. bolanderi*, blue; *I. echinospora*, brown) and hexaploid *I. occidentalis* (green) are separated as the three main populations. The hybrid species (*I.* × *herb-wagneri*, *I. maritima*, *I. sp. nov.* A, and *I. sp. nov.* B) are constructed of proportions of those three main populations.

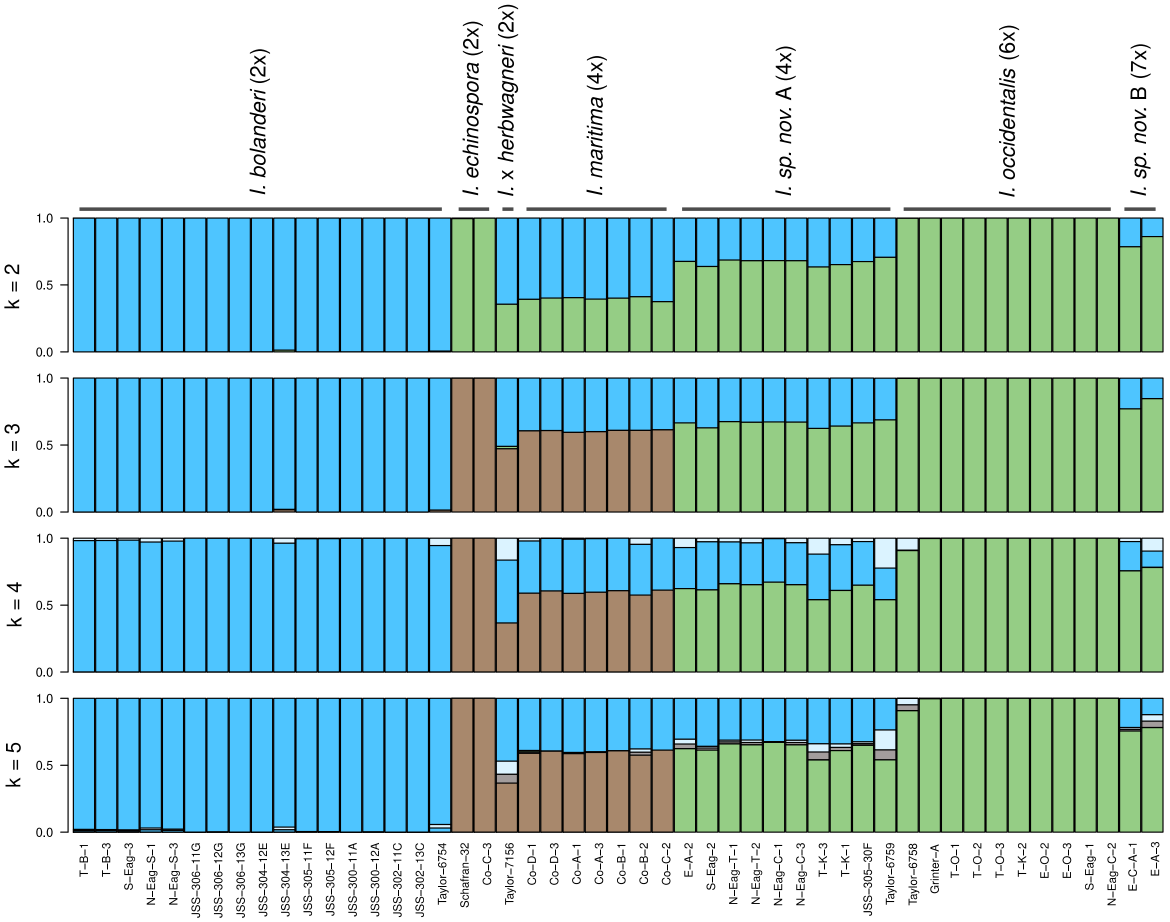

**Fig. S2** Split network depicting reticulate relationships of western North American *Isoëtes*. Each sample is colored by their taxon name. *I. echinospora* (brown); *I. sp. nov.* B (purple); *I. occidentalis* (green); *I. sp. nov.* A (red); *I. maritima* (dark blue); *I.* ✕ *herb-wagneri* (yellow); *I. bolanderi* (cyan). Hybrids of *I. echinospora* and *I. bolanderi* cluster between the two parent taxa. Hybrids between *I. occidentalis* and *I. bolanderi* also cluster between their parental taxa.

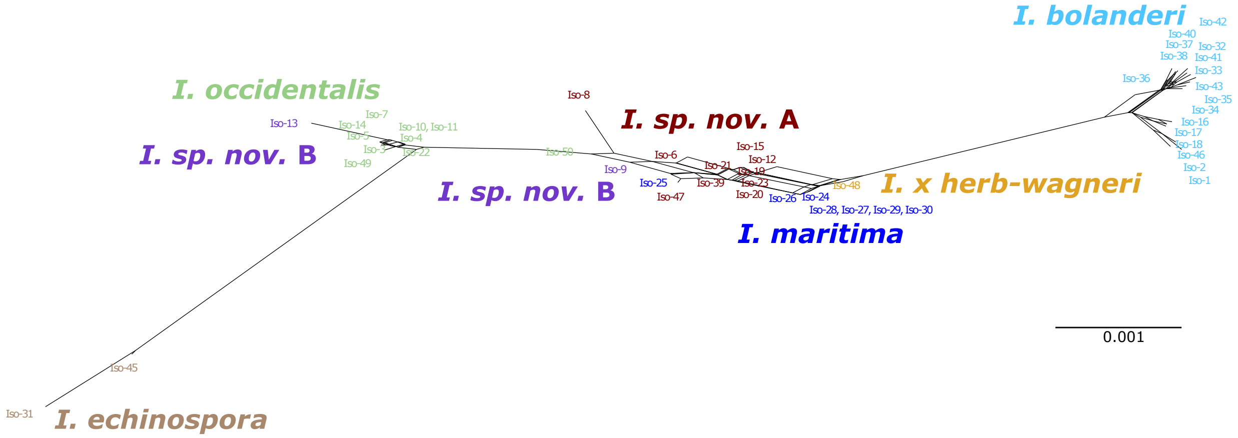


**Fig. S3** Phylogenetic tree derived from Bayesian inference (MrBayes), of North American *Isoëtes* reconstructed using multiple alleles of the *LFY* gene. Outgroups are colored in black, taxa of interest are colored in blue.

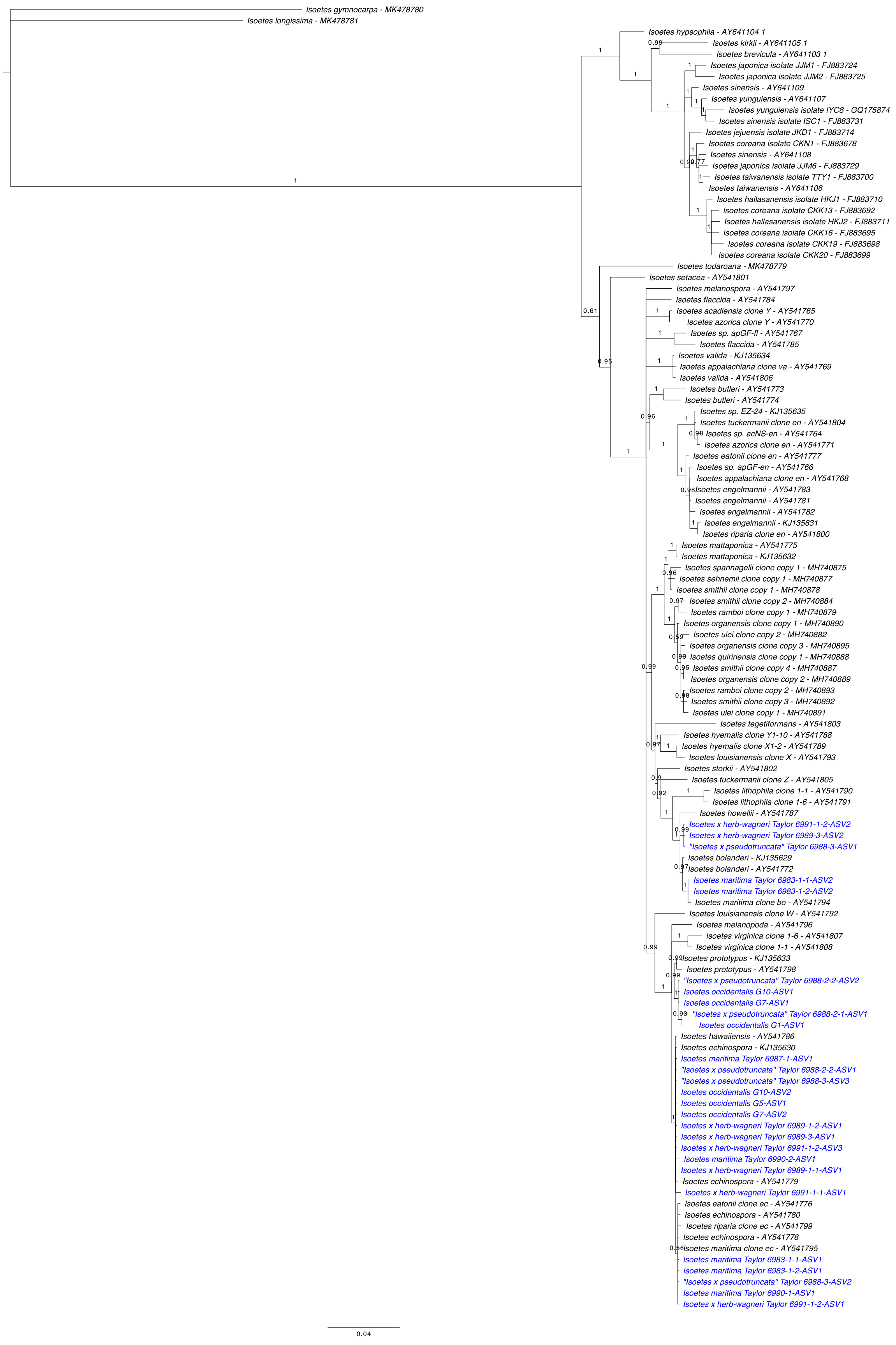
Isoetes gymnocarpa - MK478780
Isoetes longissima - MK478781
Isoetes hypsophila - AY641104 1
Isoetes kirkii - AY641105 1
Isoetes brevicula - AY641103 1
Isoetes japonica isolate JJM1 - FJ883724
Isoetes japonica isolate JJM2 - FJ883725
Isoetes sinensis - AY641109
Isoetes yunguiensis - AY641107
Isoetes yunguiensis isolate IYC8 - GQ175874
Isoetes sinensis isolate ISC1 - FJ883731
Isoetes jejuensis isolate JKD1 - FJ883714
Isoetes coreana isolate CKN1 - FJ883678
Isoetes sinensis - AY641108
Isoetes japonica isolate JJM6 - FJ883729
Isoetes taiwanensis isolate TTY1 - FJ883700
Isoetes taiwanensis - AY641106
Isoetes hallasanensis isolate HKJ1 - FJ883710
Isoetes coreana isolate CKK13 - FJ883692
Isoetes hallasanensis isolate HKJ2 - FJ883711
Isoetes coreana isolate CKK16 - FJ883695
Isoetes coreana isolate CKK19 - FJ883698
Isoetes coreana isolate CKK20 - FJ883699
Isoetes todaroana - MK478779
Isoetes setacea - AY541801
Isoetes melanospora - AY541797
Isoetes flaccida - AY541784
Isoetes acadiensis clone Y - AY541765
Isoetes azorica clone Y - AY541770
Isoetes sp. apGF-fl - AY541767
Isoetes flaccida - AY541785
Isoetes valida - KJ135634
Isoetes appalachiana clone va - AY541769
Isoetes valida - AY541806
Isoetes butleri - AY541773
Isoetes butleri - AY541774
Isoetes sp. EZ-24 - KJ135635
Isoetes tuckermanii clone en - AY541804
Isoetes sp. acNS-en - AY541764
Isoetes azorica clone en - AY541771
Isoetes eatonii clone en - AY541777
Isoetes sp. apGF-en - AY541766
Isoetes appalachiana clone en - AY541768
Isoetes engelmannii - AY541783
Isoetes engelmannii - AY541781
Isoetes engelmannii - AY541782
Isoetes engelmannii - KJ135631
Isoetes riparia clone en - AY541800
Isoetes mattaponica - AY541775
Isoetes mattaponica - KJ135632
Isoetes spannagelii clone copy 1 - MH740875
Isoetes sehnemii clone copy 1 - MH740877
Isoetes smithii clone copy 1 - MH740878
Isoetes smithii clone copy 2 - MH740884
Isoetes ramboi clone copy 1 - MH740879
Isoetes organensis clone copy 1 - MH740890
Isoetes ulei clone copy 2 - MH740882
Isoetes organensis clone copy 3 - MH740895
Isoetes quiririensis clone copy 1 - MH740888
Isoetes smithii clone copy 4 - MH740887
Isoetes organensis clone copy 2 - MH740889
Isoetes ramboi clone copy 2 - MH740893
Isoetes smithii clone copy 3 - MH740892
Isoetes ulei clone copy 1 - MH740891
Isoetes tegetiformans - AY541803
Isoetes hyemalis clone Y1-10 - AY541788
Isoetes hyemalis clone X1-2 - AY541789
Isoetes louisianensis clone X - AY541793
Isoetes storkii - AY541802
Isoetes tuckermanii clone Z - AY541805
Isoetes lithophila clone 1-1 - AY541790
Isoetes lithophila clone 1-6 - AY541791
Isoetes howellii - AY541787
Isoetes x herb-wagneri Taylor 6991-1-2-ASV2
Isoetes x herb-wagneri Taylor 6989-3-ASV2
"Isoetes x pseudotruncata" Taylor 6988-3-ASV1
Isoetes bolanderi - KJ135629
Isoetes bolanderi - AY541772
Isoetes maritima Taylor 6983-1-1-ASV2
Isoetes maritima Taylor 6983-1-2-ASV2
Isoetes maritima clone bo - AY541794
Isoetes louisianensis clone W - AY541792
Isoetes melanopoda - AY541796
Isoetes virginica clone 1-6 - AY541807
Isoetes virginica clone 1-1 - AY541808
Isoetes prototypus - KJ135633
Isoetes prototypus - AY541798
"Isoetes x pseudotruncata" Taylor 6988-2-2-ASV2
Isoetes occidentalis G10-ASV1
Isoetes occidentalis G7-ASV1
"Isoetes x pseudotruncata" Taylor 6988-2-1-ASV1
Isoetes occidentalis G1-ASV1
Isoetes hawaiiensis - AY541786
Isoetes echinospora - KJ135630
Isoetes maritima Taylor 6987-1-ASV1
"Isoetes x pseudotruncata" Taylor 6988-2-2-ASV1
"Isoetes x pseudotruncata" Taylor 6988-3-ASV3
Isoetes occidentalis G10-ASV2
Isoetes occidentalis G5-ASV1
Isoetes occidentalis G7-ASV2
Isoetes x herb-wagneri Taylor 6989-1-2-ASV1
Isoetes x herb-wagneri Taylor 6989-3-ASV1
Isoetes x herb-wagneri Taylor 6991-1-2-ASV3
Isoetes maritima Taylor 6990-2-ASV1
Isoetes x herb-wagneri Taylor 6989-1-1-ASV1
Isoetes echinospora - AY541779
Isoetes x herb-wagneri Taylor 6991-1-1-ASV1
Isoetes eatonii clone ec - AY541776
Isoetes echinospora - AY541780
Isoetes riparia clone ec - AY541799
Isoetes echinospora - AY541778
Isoetes maritima clone ec - AY541795
Isoetes maritima Taylor 6983-1-1-ASV1
Isoetes maritima Taylor 6983-1-2-ASV1
"Isoetes x pseudotruncata" Taylor 6988-3-ASV2
Isoetes maritima Taylor 6990-1-ASV1
Isoetes x herb-wagneri Taylor 6991-1-2-ASV1
0.04

**Table. S1**. This table includes the individuals used in the RADseq data, the collection numbers, herbarium they are housed in, the specimen ID, C-values (or chromosome number), and inferred ploidy levels.
